# UFold: Fast and Accurate RNA Secondary Structure Prediction with Deep Learning

**DOI:** 10.1101/2020.08.17.254896

**Authors:** Laiyi Fu, Yingxin Cao, Jie Wu, Qinke Peng, Qing Nie, Xiaohui Xie

## Abstract

**Motivation:** For many RNA molecules, the secondary structure is essential for the correct function of the RNA. Predicting RNA secondary structure from nucleotide sequences is a long-standing problem in genomics, but the prediction performance has reached a plateau over time. Traditional RNA secondary structure prediction algorithms are primarily based on thermodynamic models through free energy minimization, which imposes strong prior assumptions and is slow to run.

**Results:** Here we propose a deep learning-based method, called UFold, for RNA secondary structure prediction, trained directly on annotated data without any thermodynamic assumptions. UFold improves substantially upon previous models, with approximately 10~30% improvement over traditional thermodynamic models and 14% improvement over other learning-based methods. It achieves an F1 score of 0.91 on base pair prediction accuracy on an RNA structure prediction benchmark dataset. UFold is also fast with an inference time about 160ms per sequence up to 1600bp length. We provide an online web server that implements UFold for RNA structure prediction and is made freely available.

**Availability:** An online web server running UFold is available at https://ufold.ics.uci.edu. Code is available at https://github.com/uci-cbcl/UFold.

## 1 Introduction

The biology of RNA is diverse and complex. Aside from its conventional role as an intermediate between DNA and protein, cellular RNA consists of many other functional classes, including ribosomal RNA (rRNA), transfer RNA (tRNA), small nuclear RNA (snRNA), microRNA, and other noncoding RNAs (Noller, 1984; Rich and RajBhandary, 1976; Allmang *et al.*, 1999; Geisler and Coller, 2013). Some RNAs possess catalytic functionality, playing a role similar to protein enzymes. The spliceosome, which performs intron splicing, is assembled from several snRNAs. The microRNAs are abundant in many mammalian cell types, targeting approximately 60% of genes (Gebert and MacRae, 2019), and are often regarded as biomarkers for diverse diseases(Fu and Peng, 2017).

Cellular RNA is typically single-stranded. RNA folding is in large part determined by nucleotide base pairing, including canonical base pairing-A-U and C-G, and non-canonical base pairing-primarily G-U pairing (Fallmann *et al.*, 2017; Westhof and Fritsch, 2000). The base-paired structure is often referred to as the secondary structure of RNA(Fox and Woese, 1975). For many RNA molecules, the secondary structure is essential for the correct function of the RNA, in many cases, more than the primary sequence itself. As an evidence of this, many homologous RNA species demonstrate conserved secondary structures, accompanied by diverged sequences characterized by compensatory mutations(Mathews *et al.*, 2010).

RNA secondary structure can be determined from atomic coordinates obtained from X-ray crystallography, nuclear magnetic resonance (NMR), or cryogenic electron microscopy (Fürtig *et al.*, 2003; Cheong *et al.*, 2004; Fica and Nagai, 2017). However, these methods have low throughput. Only a tiny fraction of RNAs have experimentally determined structures. To address this limitation, experimental methods have been proposed to infer base paring by using probes based on enzymes, chemicals, and cross-linking techniques coupled with high throughput sequencing (Bevilacqua *et al.*, 2016; Underwood *et al.*, 2010). Although promising, these methods are still at the early stage of development, unable to provide precise base-pairing at a single nucleotide solution.

Computationally predicting the secondary structure of RNA is a long-standing problem in genomics and bioinformatics. Many methods have been proposed over the past two decades. They can be broadly classified into two categories: 1) single sequence prediction methods, and 2) comparative methods. In the first category, the most common method is to search for thermodynamically stable states through free energy minimization. If the secondary structure contains only nested base pairing, the energy minimization can be efficiently solved through dynamic programming, such as those implemented in Vienna RNAfold (Lorenz *et al.*, 2011), MFold (Zuker, 2003), RNAstructure (Mathews and Turner, 2006), and CONTRAfold (Do *et al.*, 2006). Faster implementations that try to improve the speed of dynamic programming include Rfold (Kiryu *et al.*, 2008), Vienna RNAplfold (Bernhart *et al.*, 2006) and LocalFold (Lange *et al.*, 2012). Efficient dynamic programming algorithms that sample suboptimal secondary structures from the Boltzmann ensembles of structures have also been proposed, e.g., CentroidFold (Sato *et al.*, 2009). However, dynamic programming breaks down when base pairs contain nonnested patterns, called pseudoknots, which includes two stem-loop structures with half of one stem intercalating between the two halves of another stem. Predicting secondary structures with pseudoknots is hard and has shown to be NP-complete under the energy minimization framework. Methods in the secondary category utilize covariance methods by aligning related RNA sequences and identifying correlated compensatory mutations. Although the list of proposed methods in each of two categories is long and diverse (Kings Oluoch *et al.*, 2018), the performance of these methods has not been significantly improved over time, reaching a performance ceiling of about 80% (Seetin and Mathews, 2012). It is possibly because they fail to account for base pairing resulting from tertiary interactions (Nowakowski and Tinoco Jr, 1997), unstacked base pairs, pseudoknot, noncanonical base pairing, or other unknown factors (Westhof and Fritsch, 2000).

Recently deep learning techniques have started to emerge as an alternative approach to the functional structure prediction problems including RNA secondary structure prediction problem (Zhang *et al.*, 2019; Wang *et al.*, 2019; Chen *et al.*, 2019a; Singh *et al.*, 2019; Wang *et al.*, 2016). Compared to the thermodynamic model-based approaches, the learning-based methods benefits from making few assumptions, allowing pseudoknots, and accounting for tertiary interactions, noncanonical base pairing, or other previously unrecognized base pairing constraints. Existing deep learning methods differ in model architectural design and their choices of model input and output. These methods either treat the input as a sequence, utilizing LSTM (Hochreiter and Schmidhuber, 1997) or transformer encoder (Cer *et al.*, 2018) to capture long-range interactions between nucleotides (Sato *et al.*, 2020; Singh *et al.*, 2019; Chen *et al.*, 2019b). Other methods aim to integrate deep learning techniques with dynamic programming or thermodynamic methods to alleviate prediction biases (Wang *et al.*, 2019; Zhang *et al.*, 2019; Sato *et al.*, 2020). However, existing deep learning approaches still face several challenges: First, both LSTM and transformer encoder modules involve a huge number of model parameters, which lead to high computational cost and low efficiency. Second, integrating with thermodynamic optimization methods will push the models to assume the assumptions underlying traditional methods, which can hinder the model performance. Third, because the performance of deep learning models depends heavily on the distribution of training data, we need to think about how to improve the performance of these models on previously unseen classes of RNA structures (Sato *et al.*, 2020).

Instead of using the nucleotide sequence itself, the input of our model consists of all possible base-pairing maps within the input sequence. Each map, first represented by a square matrix of the same dimension as the input sequence length, denotes the occurrences of one of the 16 possible base pairs between the input nucleotides. Under this new representation, the input is treated as a 2D “image” with 16 channels, allowing the model to explicitly consider all long-range interactions and all possible base pairing, including non-canonical ones. We include one additional channel to store the pairing probability between input base pairs calculated based on three paring rules (Zhang *et al.*, 2019) and concatenate it with the previous 16 channel representation. So an overall 17 channel 2D map is used as our model input. We use an encoder-decoder framework to extract multi-scale long- and short-range interaction features of the input sequence, implemented in a U-Net model. For this reason, we will refer to our method as UFold (stands for U-Net based on RNA folding). The output of UFold is the predicted contact score map between the bases of the input sequence. UFold is fully convolutional, and as such it can readily handle input sequences with variable length.

We conduct experiments on both known family RNA sequences and cross family RNA sequences to compare the performance of UFold against both the traditional energy minimization-based methods and recent learning-based methods. We show that UFold yields substantial performance gain over previous methods, highlighting its promising potential in solving the RNA secondary structure prediction problem.

UFold is fast with an inference time of average 160 ms per sequence for RNA sequences with lengths of up to 1600bp. We have developed an online web server running UFold RNA secondary structure prediction. The server is freely available, allowing users to enter sequences and visualize predicted secondary structures.

## 1 Methods

### 2.1 Datasets

Several benchmark datasets are used in this study: a) RNAStralign (Tan *et al.*, 2017), which contains 30,451 unique sequences from 8 RNA families; b) ArchiveII (Sloma and Mathews, 2016), which contains 3,975 sequences from 10 RNA families and is the most widely used dataset for benchmarking RNA structure prediction performance; c) bpRNA-1m (Danaee *et al.*, 2018), which contains 102,318 sequences from 2,588 families and is one of the most comprehensive RNA structure datasets available; and d) bpRNA-new, derived from Rfam 14.2 (Kalvari *et al.*, 2021), containing sequences from 1,500 new RNA families. RNA families occurring in bpRNA-1m or any other dataset are excluded from bpRNA-new. In this work, bpRNA-new dataset is treated as a cross-family dataset to assess cross-family model generalization.

The RNAStralign dataset is randomly split into training, validation and test sets, with 24,895, 2,702, and 2,854 samples, respectively. Redundant sequences between test and training are removed. For bpRNA-1m dataset, we follow the same process procedure used in SPOT-RNA (Singh *et al.*, 2019) by using CD-HIT program (Li and Godzik, 2006) to remove redundant sequences and randomly split the dataset into three sub-datasets for training and testing, called TR0 and TS0, respectively. Redundancy removed ArchiveII and bpRNA-new are used only for testing.

### 2.2 Input and output representation

The general problem of the RNA secondary structure prediction is to predict base pairing patterns given an input sequence. Let ***x*** = (*x*_1_, *x*_2_,…,*x*_*L*_) with *x*_*i*_ ∈ {′A′, ′U′, ′C′, ′G′} be an input sequence of length *L*. The goal is to predict the secondary structure of *x*, represented by a contact matrix *A* ∈ {0,1}^*L*×*L*^ with *A*_*ij*_ = 1 denoting a base pairing between bases *x*_*i*_ and *x*_*j*_, and 0 otherwise. UFold utilizes a deep neural network to predict the contact matrix given the input. Next, we describe several design choices behind UFold (**Fig. 1**).

**Fig. 1.**
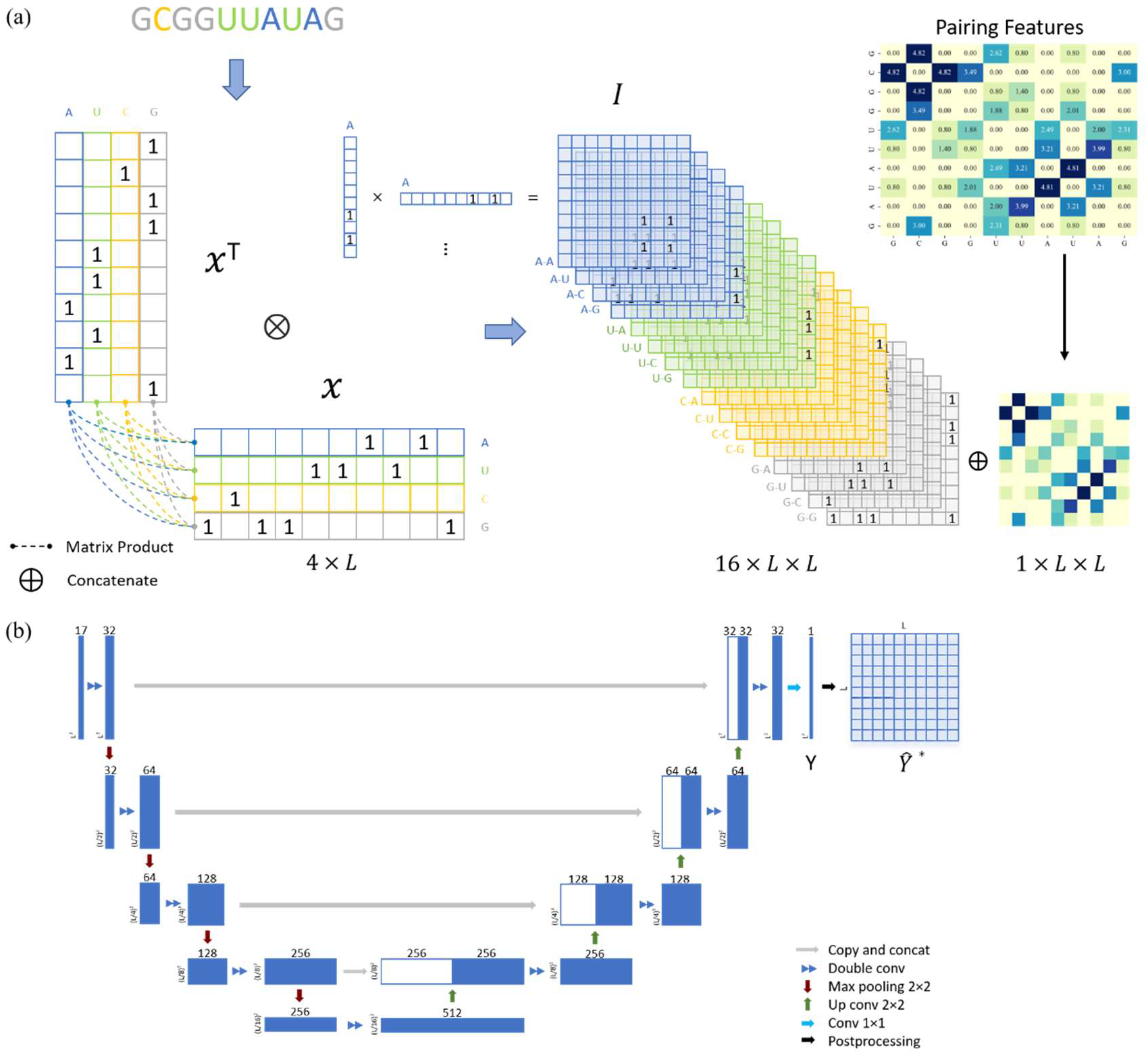
The overall architecture of UFold. (a) The input sequence is first converted into one-hot representation. A novel representation of the sequence is then introduced by taking outer product of all combinations of base pair channels, resulting in an image-like representation with 16 channels and with the same size as the contact map. We calculate a paring possibilities matrix according to three paring rules and concatenate this extra matrix with previous feature to obtain the final 17 channel input. (b) Detailed architecture of our framework. The input is a 17 × *L* × *L* tensor representation of the original sequence. The U-Net takes the 17 × *L* × *L* tensor as input and outputs an *L* × *L* symmetric score matrix Y. After postprocessing, matrix *Ŷ*\* is the final prediction of the contact map.

Most existing learning-based methods treat the input as a sequence and use recurrent neural nets (RNNs) to model the interaction between different bases. Gated RNNs, such as LSTMs and GRUs, are often the method of choice for dealing with sequential data because of their ability to model long range dependencies. However, RNN models need to be run sequentially, causing issues in both training and inference. Newer RNA structure prediction models based on transformers, which does not require the sequential data to be processed in order, have also been proposed.

Unlike the previous models, UFold converts the input sequence into an “image”. This is done by first encoding *x* with one-hot representation, representing the sequence with an *L* × 4 binary matrix *X* ∈ {0,1}^×4^. ***x*** is then transformed into a 16 × *L* × *L* tensor through a Kronecker product between ***x*** and itself, followed by reshaping dimensions (**Fig. 1a**),

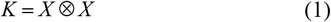

In this representation, input *K* ∈{0,1}^16×*L*×*L*^ can be understood as an image of size *L* × *L* with 16 color channels. Each channel specifies one of the 16 possible base pairing rules; *K*(*i*, *j*, *k*) denotes whether bases *x*_*j*_ and *x*_*k*_ are paired according to the *i-*th base pairing rule (e.g., i=2 for A-C pairing).

To overcome the sparsity bringing by converting sequencing into 16 channels, we also adopt an extra channel used in CDPFold (Zhang *et al.*, 2019), which reflects the implicit matching between bases. Basically, we calculate the paring possibilities between each nucleotide and others from one sequence according to three paring rules (Zhang *et al.*, 2019), using these rules we could calculate the specific values of each nucleotide position with other nucleotides. These non-binary values may help alleviate the sparsity of the model and provide more information on paring bases. The calculated matrix *W* ∈ ℝ^1×*L*×*L*^ is then concatenated with *K* along the first dimension to get the final UFold input *I* of dimension 17 × *L* × *L*.

UFold takes *I* as input and computes *Y* = *f*(*I*; *θ*) with a deep convolutional neural net (**Fig. 1b**). The output *Y* ∈ [0,1]^*L* × *L*^ is an *L* × *L* matrix, with *Y*_*ij*_ denoting the probability score of nucleotides bases *x*_*i*_ and *x*_*j*_ being paired.

The new input representation taken by UFold has several advantages: First, using an image representation allows it to model all possible longrange interactions explicitly. Base pairing between distant sequence segments shows up locally in the image representation. Second, it considers all possible base pairing patterns, making no distinction between canonical and non-canonical base pairs. Third, it allows us to implement a fully convolutional neural model that can handle variable sequence length, eliminating the need of padding the input sequence to a fixed length.

### 2.3 Input and Scoring network architecture

UFold uses an encoder-decoder architecture for computing predicted contact score matrix *Y* (**Fig. 1**). The model consists of a sequence of downsampling layers (encoder) to derive increasingly complex semantic representations of the input, followed by a sequence of up-sampling layers (decoder), with lateral connections from the encoder to fill in contextual information. The overall design follows the U-Net model, widely used in the field of image segmentation.

All operations in UFold are fully convolutional. Thus, the input sequence can be of variable length, with the output matrix changing correspondingly. This feature is especially beneficial for RNA secondary structure as the range of the input sequence length is very large, from tens of nucleotides for small RNAs to thousands of nucleotides for large RNAs. Padding input sequences to the same length as done in other methods would have significantly impacted the efficiency of the algorithm.

UFold is trained by minimizing the cross-entropy between the predicted probability contact matrix *Y* and the true contact matrix *A*, using stochastic gradient descent. A positive weight *ω* of 300 is added to leverage the imbalanced 0/1 distribution to derive the loss function as below,

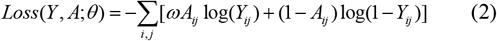

### 2.4 Postprocessing

After the symmetric contact scoring matrix *Y* is computed by UFold, we use a postprocessing procedure to derive the final secondary structure. The postprocessing procedure takes into account four hard constraints in the secondary structure: 1) the contact matrix should be symmetric; 2) only canonical plus U-G paring rules are allowed (this can be relaxed by including other non-canonical base pairs); 3) no sharp loops are allowed, for which we set *A*_*ij*_ = 0, ∀*i*, *j* with |*i* − *j*| < 4; and 4) no overlapping pairs are allowed, that is, *A***1** ≤ **1**. We follow the steps used in E2Efold by encoding constraints 2 & 3 into a matrix *M*, defined as *M*(*x*)_*ij*_ ≔ 1 if nucleotides *x* and *x* can be paired under constraints 2 & 3 and equals to 0 otherwise.

To address the first two constraints, we transform Y according to

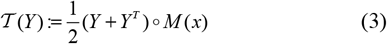

where ° denotes element-wise multiplication. It ensures that the transformed Y is symmetric and satisfies constraints 1, 2 and 3.

To address the last constraint, we relax it into a linear programming problem,

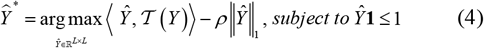

which tries to find an optimal scoring matrix *Ŷ* that is most similar to 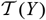 while at the same time satisfying the nonoverlapping pair constraint. The similarity is measured in terms of the inner product between *Ŷ* and 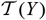. *ρ* is a hyperparameter controlling the sparsity of the final output.

The final predicted binary contact map is taken to be *Y* ∗ after thresholding it with an offset, which is chosen through a grid search.

### 2.5 Training and Evaluation

During training, stratified sampling (Chen et al., 2019a) is applied to the training set to balance the number of training samples from each RNA family. The hyperparameters of UFold are tuned based on the validation set.

To improve model transferability on previously unknown RNA families, we augment the training set with synthetic data to train UFold. The synthetic data are generated by randomly mutating sequences in the bpRNA-new dataset (previously unseen RNA families). We then use RNAfold to generate predicted structures on the synthetic data and treat them as ground-truth.

For evaluation, we report three metrics: F1 score, precision and recall. Precision is defined as 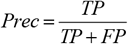, evaluated on all predicted base pairs. Recall is defined as 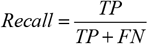. And And F1 score is the harmonic mean of precision and recall, defined as 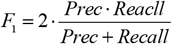.

## 3 Results

To benchmark the performance of different models, we conduct two experimental studies: A) train models on the RNAStralign training set and evaluate on the RNStralign test set and ArchiveII; and B) train models on the bpRNA-1m training set (TR0) and evaluate on the bpRNA-1m test set (TS0) as well as on bpRNA-new. Published deep learning models usually report results from either Study A or Study B. To have a fair and direct comparison with previous models, we report results from both, following the same data splitting, preprocessing, and evaluation protocols.

In comparing the results from different models, we treat within- vs cross-family results separately. In both studies, the test sets, except bpRNA-new, contain mostly within family RNA species, i.e., RNA species from a similar family occurring in the training set. By contrast, the bpRNA-new dataset contains only cross-family RNA species, that is, none of them shares the same RNA family as those in the training set. Although most RNAs are from a known family, it is necessary to consider the model’s performance on previously unseen families to assess its model transferability.

### 3.1 Experimental results on within family datasets

In this section, we report the results of our model on within-family test sets. **Table 1** summarizes the evaluation results of UFold on the ArchieveII test set (from Study A), together with the results of a collection of traditional energy-based and recent learning-based methods (Chen *et al.*, 2019a). The traditional methods achieve an F1 score in the range of 0.42 to 0.61. A recent state-of-the-art learning-based method improves the F1 score to 0.80 (MXfold2). UFold can further improve the performance, achieving an F1 score of 0.91. Compared with MXfold2, UFold achieves a 14% increase in F1 score, a 19% increase in recall, and a 7.5% increase in precision.

**Table 1.** Benchmark results on the ArchiveII dataset.

| Method | Prec | Rec | F1 |
| --- | --- | --- | --- |
| UFold | **0.887** | **0.928** | **0.905** |
| MXfold2 | 0.825 | 0.780 | 0.796 |
| SPOT-RNA | 0.743 | 0.726 | 0.711 |
| E2Efold | 0.734 | 0.660 | 0.686 |
| CDPfold | 0.557 | 0.535 | 0.545 |
| LinearFold | 0.641 | 0.617 | 0.621 |
| MFold | 0.428 | 0.383 | 0.401 |
| RNAstructure | 0.563 | 0.615 | 0.585 |
| RNAfold | 0.565 | 0.627 | 0.592 |
| CONTRAFold | 0.607 | 0.679 | 0.638 |

**Table 2** summarizes the evaluation results on the TS0 test set (from Study B). Since this dataset was also used in two other deep learning-based methods - SPOT-RNA and MXfold2, we compare UFold with these two methods along with two energy-based methods. Again, UFold outperforms both the deep learning-based and the energy-based methods. UFold achieves an F1 score of 0.654 on this dataset, corresponding to a 5.6% improvement over SPOT-RNA, the state-of-the-art method on this dataset, and 13.7% improvement over traditional methods. Improvements in recall and precision also surpass all other methods.

**Table 2.** Benchmark results on the TS0 dataset.

| Method | Prec | Rec | F1 |
| --- | --- | --- | --- |
| UFold | **0.606** | **0.741** | **0.654** |
| SPOT-RNA | 0.594 | 0.693 | 0.619 |
| MXfold2 | 0.519 | 0.646 | 0.558 |
| ContextFold | 0.583 | 0.597 | 0.575 |
| RNAfold | 0.446 | 0.631 | 0.508 |

Predicting secondary structures with pseudoknots is especially challenging for thermodynamic models. We also validate the performance of UFold on predicting base pairing in the presence of pseudoknots. For this purpose, we pull out all RNA structures with pseudoknots in the RNAS-tralign test set, on which we then benchmark UFold against two other methods that can predict pseudoknots, including SPOT-RNA, E2Efold and RNAstructure. As shown in **Table 3**, all other methods tend to misclassify normal structures as pseudoknots. By contrast, UFold still achieves much less false positive (FP) predictions (49 v. 591/242/307) while maintains high sensitivity (TP 1247 v. 1237/1312/1248), highlighting the robustness of UFold in the presence of pseudoknots.

**Table 3.** Evaluation results of RNA structures with pseudoknots on the RNAStralign test dataset.

| Method | TP | FP | TN | FN |
| --- | --- | --- | --- | --- |
| UFold | **1247** | **49** | **1491** | **3** |
| SPOT-RNA | 1237 | 591 | 35 | 959 |
| E2Efold | 1312 | 242 | 1271 | 0 |
| RNAstructure | 1248 | 307 | 983 | 286 |

### 3.2 Experimental results on cross family datasets

In this section, we evaluate the performance of UFold on previously unseen RNA families. We expect learning-based methods do poorly on these RNAs since they are not represented in the training set. To address this problem, methods integrating free energy minimization with deep learning methods have been proposed, like MXfold2 (Sato *et al.*, 2020). However, these methods inadvertently introduce biases into the prediction model and likely lead to reduced performance on within family RNAs.

UFold does not involve any energy minimization term in its model. Instead, it uses data augmentation to improve the performance on cross-family RNAs. 10,000 synthetic RNA sequences are generated by randomly mutating real RNA sequences. All the synthetic sequences have passed CD-HIT 80. Secondary structures predicted from an energy-based method (RNAfold was used) are treated as the ground-truth and are merged with the TR0 training set for model training.

**Table 4** shows the evaluation results of UFold on the bpRNA-new dataset, containing about 1,500 previously unseen RNA families. UFold can achieve a similar performance as other methods like MXfold2, all of which involve thermodynamic terms or constraints in their objectives. By contrast, UFold is a pure learning-based method. Through data augmentation, it can learn to predict the structures of RNAs not represented in the training set.

**Table 4.** Benchmark results on the bpRNA-new dataset.

| Method | Prec | Rec | F1 |
| --- | --- | --- | --- |
| UFold | 0.537 | **0.724** | 0.608 |
| MXfold2 | **0.585** | 0.710 | **0.632** |
| SPOT-RNA | 0.599 | 0.619 | 0.596 |
| ContextFold | 0.595 | 0.539 | 0.554 |
| RNAfold | 0.552 | 0.720 | 0.617 |

### 3.3 Visualization

After quantitively evaluating the prediction performance, we visualize the RNA secondary structures predicted by UFold to check the pairing details of each nucleotide. For this purpose, the predicted contact maps were first converted to a dot-bracket format according to base pair positions. Raw sequences with the corresponding predicted dot-bracket sequences were fed into the ViennaRNA web server powered by Forna (Kerpedjiev *et al.*, 2015) to obtain the visualization result. As a comparison, we also show the predicted structures from other three best performed methods, MXfold2, SPOT-RNA and E2Efold as well as the ground-truth structures. Two examples from arp_Aspe.fumi._GSP-41122.ct and 16S_rRNA/Alphaproteobacteria/L41814.ct respectively are drawn and shown in **Fig. 2**. In both cases, UFold generates RNA secondary structures more similar to the ground-truth when compared with other state-of-the-art methods like MXfold2, SPOT-RNA and E2Efold, showing the closest secondary structure to the ground truth structure.

**Fig. 2.**
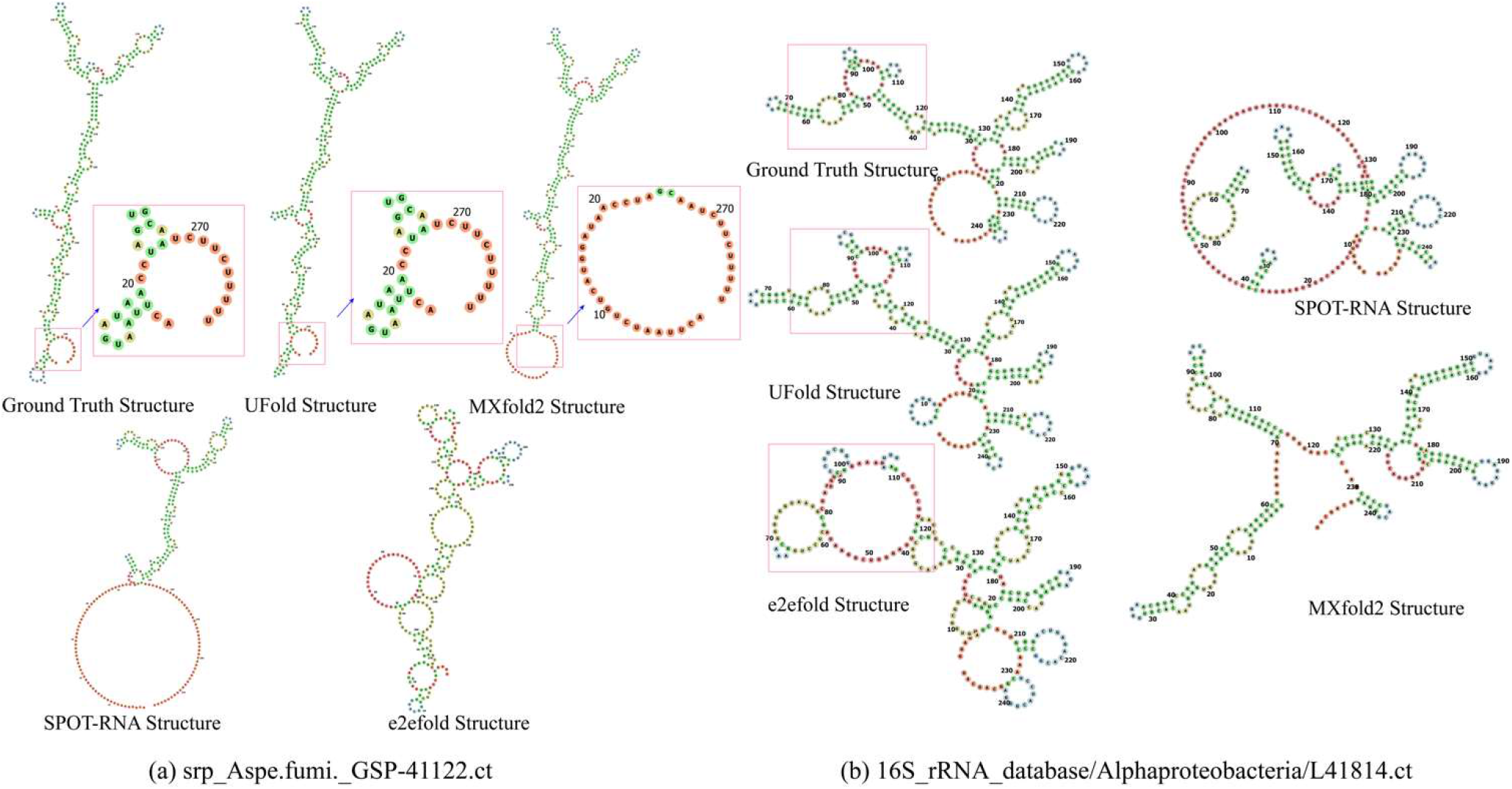
Visualization of two example UFold RNA secondary structure predictions. From top to bottom: ground truth, UFold prediction, and E2Efold prediction. Two RNA sequences are (a) arp_Aspe.fumi._GSP-41122.ct and (b) 16S_rRNA/Alphaproteobacteria/L41814.ct. Major prediction differences are highlighted by red boxes. In both cases, UFold produces predictions more aligned with the ground-truth.

As a test case, we applied UFold to predict secondary structures of SARS-CoV-2 RNA sequences, the coronavirus causing the COVID-19 pandemic(Huston *et al.*, 2020; Zhou *et al.*, 2020). SARS-CoV-2 is roughly 30 kilobases long, 97% of which have not been structurally explored. We used UFold to predict some of the key structures extracted from its original sequence and compared the predicted structures with experimentally determined structures published recently (Wacker et al., 2020). As shown in **Fig. 3**, our predicted structures show high consistencies with the experimentally determined structures. Because SARS-CoV-2 is a new virus not contained in any of the training datasets, the good consistencies show the effectiveness of UFold. Moreover, we can generate predicted RNA structures for all regions of the virus.

**Fig. 3.**
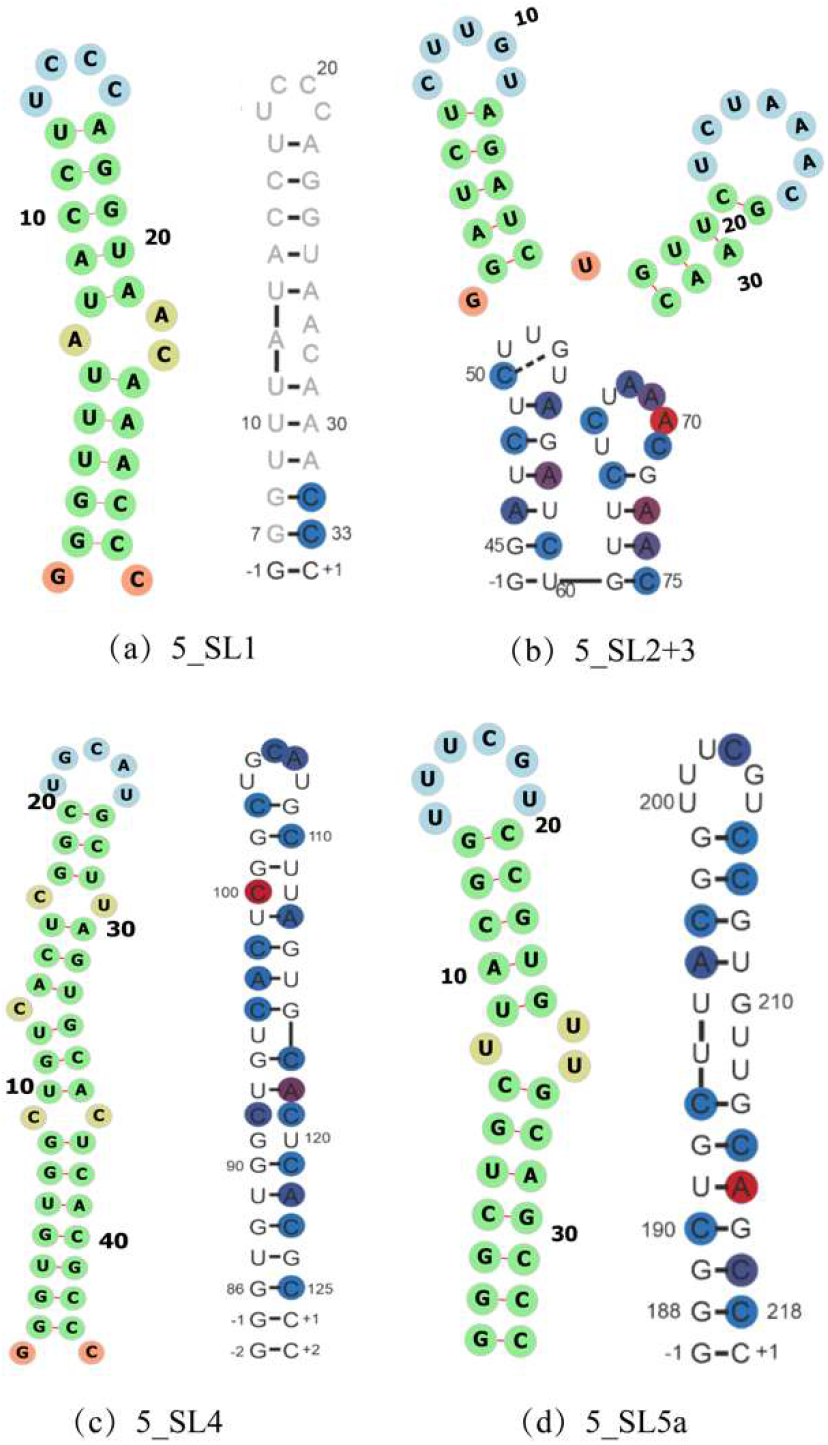
SARS-Cov2 virus RNA secondary structure. Four different regions of the virus are shown (a-d). RNA secondary structures predicted by UFold are shown on the left (a, c, d) or top (b) with paired bases colored green. NMR-DMS experimentally determined structures are shown on the right (a, c, d) or the bottom (b) with paired bases colored blue or red. It shows a good consistency between UFold predicted and experimentally determined secondary structures.

### 3.4 Inference time

The speed of the prediction algorithm is an important factor in RNA secondary structure prediction, especially for multiple sequences predicting simultaneously. Traditional energy minimization-based methods tend to be slow because of the time complexity of the minimization algorithm. Deep learning-based methods like MXfold2 and SPOT-RNA utilize LSTM structure, which require significantly more parameters than UFold, resulting in low efficiency. UFold inference, on the other hand, runs on feedforward neural nets only. Specifically, it comprises of fully connected convolutional neural network, which greatly reduces the running time since all operations are readily parallelizable. It can also handle multiple sequences at once, leading to significantly higher throughput.

The average inference time per sequence of UFold on the RNAStralign test set (containing sequences longer than 1000bp) is reported in **Table 5**, together with the average running times of other methods. UFold is much faster than both learning-based and energy-based methods. UFold is nearly two-times faster than MXfold2, and orders-of-magnitude faster than RNAstruture, a popular energy-based method that can handle pseudoknots. The running times of UFold and three other recent deep learning-based methods are also shown in **Table 5**. All these methods are implemented in pytorch(Paszke *et al.*, 2019) and thus it allows us to compare their model efficiency directly. Our model is still the fastest one among all the other deep learning methods, further demonstrating the efficiency of UFold.

**Table 5:** Inference time on the RNAStralign test set.

| Method | Time per seq |
| --- | --- |
| **UFold (Pytorch)** | **0.16s (GPU)** |
| MXfold2(Pytorch) | 0.31s (GPU) |
| E2Efold (Pytorch) | 0.40s (GPU) |
| SPOT-RNA(Pytorch) | 77.80s (GPU) |
| CDPfold (tensorflow) | 300.107s |
| LinearFold (C) | 0.43s |
| Mfold (C) | 7.65s |
| RNAstructure (C) | 142.02s |
| RNAfold (C) | 0.55s |
| CONTRAFold (C) | 30.58s |

### 3.5 Web server

To facilitate the accessibility of UFold, we developed a web server running UFold on the backend and made it freely available. Users can type in or upload RNA sequences in FASTA format. Our server predicts RNA secondary structures using the pre-trained UFold model and stores predicted structures in a dot-bracket file for end-users to download. The server also provides an interface connection to the ViennaRNA web service powered by forna tool (Kerpedjiev *et al.*, 2015) for visualizing predicted structures. Most existing RNA prediction servers only permit predicting one RNA sequence at a time, such as RNAfold, MXfold2, and SPOT-RNA, and restrict the length of the input sequence. Our server does not have such limitations. Its main functionality differences compared to other servers are highlighted in **Table 6**.

**Table 6.** Functionality comparison of different RNA structure prediction web servers.

| Supported Functions | Servers |  |  |  |
| --- | --- | --- | --- | --- |
|  | UFold | SPOT-RNA | RNAfold | MXfold2 |
| Sequence type-in | **Yes** | **Yes** | **Yes** | **Yes** |
| Fasta file | **Yes** | No | **Yes** | No |
| Length > 600 bp | **Yes** | **Yes** | No | No |
| Support multi-samples | **Yes** | No | No | No |

## 4 Discussion

In this study, we present UFold, a new deep learning-based model for RNA secondary structure prediction. We show that UFold outperforms previous methods by a large margin, achieving 10~30% performance improvement over traditional thermodynamic methods and 14% improvement in F1 score over the state-of-the-art learning-based method on standard dataset. UFold achieves a F1 score of 0.905 on the ArchiveII dataset and recall score of over 0.70 on bpRNA dataset, bringing in substantial gains in RNA secondary prediction accuracy. In addition, UFold is fast, being able to generate predictions at roughly 160ms per sequence.

A key difference between UFold and previous learning-based methods is its architectural design. Instead of using raw sequences as input, UFold converts sequences into “images”, explicitly modeling all possible base pairing between the nucleotides of the input sequence. This choice of input representation has several important implications: First, base pairing patterns between distant sequence segments show up locally in the image representation, making the detection and learning of these distant base pairing patterns easier. Second, all base pairing patterns are explicitly represented in the input, allowing the model to pick up all potential base pairing rules that might contribute to the formation of the secondary structure. Lastly, but perhaps most importantly, the image representation allows us to implement a fully convolutional model to pick up base pairing features across multiple scales through an encoder-decoder architecture. This implementation is not only efficient, with operations highly parallelable and allowing for variable input sequence length, but also highly effective in combining both local and global features for the final prediction.

Although UFold demonstrates a great potential in solving the RNA secondary structure prediction problem, as a learning-based method, its performance is inevitably closely attached to the quality of training data. Unfortunately, the number of experimentally resolved RNA secondary structures through X-ray crystallography or NMR remains small. Many secondary structures in the RNAStralign dataset are computationally generated by aligning homologous sequences. Fortunately, high-throughput methods for determining or constraining the secondary structures of RNAs are starting to emerge (Strobel *et al.*, 2018; Lusvarghi *et al.*, 2013). Because UFold uses a flexible network architecture, we expect it to be able to incorporate the high-throughput data to improve model training and inference.

We should note that the method presented here can potentially be applied for protein structure prediction as well. The number of amino acids is much higher than the number of bases. It is worth exploring whether all amino acid pairs, which is 400, or a subset of them should be considered in the input representation.

In summary, we show the promising potential of deep learning in solving the long-standing RNA secondary structure problem. The new framework presented here brings in a significant performance gain. We expect the prediction accuracy to be further improved as more and higher quality training data are becoming available.

## Acknowledgements

We acknowledge helpful discussions with MH Celik and members of the Xie lab.

## References

Allmang, C. et al. (1999) Functions of the exosome in rRNA, snoRNA and snRNA synthesis. The EMBO journal, 18, 5399–5410.

Bernhart, S.H. et al. (2006) Local RNA base pairing probabilities in large sequences. Bioinformatics, 22, 614–615.

Bevilacqua, P.C. et al. (2016) Genome-wide analysis of RNA secondary structure. Annual review of genetics, 50, 235–266.

Cer, D. et al. (2018) Universal sentence encoder. arXiv preprint arXiv:1803.11175.

Chen, X. et al. (2019a) RNA secondary structure prediction by learning unrolled algorithms. In, International Conference on Learning Representations.

Chen, X. et al. (2019b) RNA secondary structure prediction by learning unrolled algorithms. In, International Conference on Learning Representations.

Cheong, H.-K. et al. (2004) Rapid preparation of RNA samples for NMR spectroscopy and X-ray crystallography. Nucleic acids research, 32, e84–e84.

Danaee, P. et al. (2018) bpRNA: large-scale automated annotation and analysis of RNA secondary structure. Nucleic acids research, 46, 5381–5394.

Do, C.B. et al. (2006) CONTRAfold: RNA secondary structure prediction without physics-based models. Bioinformatics, 22, e90–e98.

Fallmann, J. et al. (2017) Recent advances in RNA folding. Journal of biotechnology, 261, 97–104.

Fica, S.M. and Nagai, K. (2017) Cryo-electron microscopy snapshots of the spliceosome: structural insights into a dynamic ribonucleoprotein machine. Nature Structural & Molecular Biology, 24, 791.

Fox, G.E. and Woese, C.R. (1975) 5S RNA secondary structure. Nature, 256, 505–507.

Fu, L. and Peng, Q. (2017) A deep ensemble model to predict miRNA-disease association. Scientific reports, 7, 1–13.

Fürtig, B. et al. (2003) NMR spectroscopy of RNA. ChemBioChem, 4, 936–962.

Gebert, L.F. and MacRae, I.J. (2019) Regulation of microRNA function in animals. Nature reviews Molecular cell biology, 20, 21–37.

Geisler, S. and Coller, J. (2013) RNA in unexpected places: long non-coding RNA functions in diverse cellular contexts. Nature reviews Molecular cell biology, 14, 699–712.

Hochreiter, S. and Schmidhuber, J. (1997) Long short-term memory. Neural computation, 9, 1735–1780.

Huston, N.C. et al. (2020) Comprehensive in-vivo secondary structure of the SARS-CoV-2 genome reveals novel regulatory motifs and mechanisms. Molecular Cell.

Kalvari, I. et al. (2021) Rfam 14: expanded coverage of metagenomic, viral and microRNA families. Nucleic Acids Research, 49, D192–D200.

Kerpedjiev, P. et al. (2015) Forna (force-directed RNA): Simple and effective online RNA secondary structure diagrams. Bioinformatics, 31, 3377–3379.

Kings Oluoch, I. et al. (2018) A Review on RNA Secondary Structure Prediction Algorithms. In, 2018 International Congress on Big Data, Deep Learning and Fighting Cyber Terrorism (IBIGDELFT). IEEE, ANKARA, Turkey, pp. 18–23.

Kiryu, H. et al. (2008) Rfold: an exact algorithm for computing local base pairing probabilities. Bioinformatics, 24, 367–373.

Lange, S.J. et al. (2012) Global or local? Predicting secondary structure and accessibility in mRNAs. Nucleic acids research, 40, 5215–5226.

Li, W. and Godzik, A. (2006) Cd-hit: a fast program for clustering and comparing large sets of protein or nucleotide sequences. Bioinformatics, 22, 1658–1659.

Lorenz, R. et al. (2011) ViennaRNA Package 2.0. Algorithms for molecular biology, 6, 26.

Lusvarghi, S. et al. (2013) RNA secondary structure prediction using high-throughput SHAPE. JoVE (Journal of Visualized Experiments), e50243.

Mathews, D.H. et al. (2010) Folding and finding RNA secondary structure. Cold Spring Harbor perspectives in biology, 2, a003665.

Mathews, D.H. and Turner, D.H. (2006) Prediction of RNA secondary structure by free energy minimization. Current opinion in structural biology, 16, 270–278.

Noller, H.F. (1984) Structure of ribosomal RNA. Annual review of biochemistry, 53, 119–162.

Nowakowski, J. and Tinoco Jr, I. (1997) RNA structure and stability. In, Seminars in virology. Elsevier, pp. 153–165.

Paszke, A. et al. (2019) PyTorch: An imperative style, high-performance deep learning library. In, Wallach, H. et al. (eds), Advances in neural information processing systems 32. Curran Associates, Inc., pp. 8024–8035.

Rich, A. and RajBhandary, U. (1976) Transfer RNA: molecular structure, sequence, and properties. Annual review of biochemistry, 45, 805–860.

Sato, K. et al. (2009) CENTROIDFOLD: a web server for RNA secondary structure prediction. Nucleic acids research, 37, W277–W280.

Sato, K. et al. (2020) RNA secondary structure prediction using deep learning with thermodynamic integration. bioRxiv.

Seetin, M.G. and Mathews, D.H. (2012) RNA structure prediction: an overview of methods. In, Bacterial Regulatory RNA. Springer, pp. 99–122.

Singh, J. et al. (2019) RNA secondary structure prediction using an ensemble of two-dimensional deep neural networks and transfer learning. Nature communications, 10, 1–13.

Sloma, M.F. and Mathews, D.H. (2016) Exact calculation of loop formation probability identifies folding motifs in RNA secondary structures. RNA, 22, 1808–1818.

Strobel, E.J. et al. (2018) High-throughput determination of RNA structures. Nat Rev Genet, 19, 615–634.

Tan, Z. et al. (2017) TurboFold II: RNA structural alignment and secondary structure prediction informed by multiple homologs. Nucleic Acids Research, 45, 11570–11581.

Underwood, J.G. et al. (2010) FragSeq: transcriptome-wide RNA structure probing using high-throughput sequencing. Nature methods, 7, 995–1001.

Wang, L. et al. (2019) DMFold: A novel method to predict RNA secondary structure with pseudoknots based on deep learning and improved base pair maximization principle. Frontiers in genetics, 10, 143.

Wang, S. et al. (2016) Protein Secondary Structure Prediction Using Deep Convolutional Neural Fields. Scientific Reports, 6, 1–11.

Westhof, E. and Fritsch, V. (2000) RNA folding: beyond Watson–Crick pairs. Structure, 8, R55–R65.

Zhang, H. et al. (2019) A new method of RNA secondary structure prediction based on convolutional neural network and dynamic programming. Frontiers in genetics, 10, 467.

Zhou, P. et al. (2020) A pneumonia outbreak associated with a new coronavirus of probable bat origin. Nature, 579, 270–273.

Zuker, M. (2003) Mfold web server for nucleic acid folding and hybridization prediction. Nucleic acids research, 31, 3406–3415.

